# Insights into Functional Connectivity in Mammalian Signal Transduction Pathways by Pairwise Comparison of Protein Interaction Partners of Critical Signaling Hubs

**DOI:** 10.1101/2019.12.30.891200

**Authors:** Chilakamarti V. Ramana

**Affiliations:** Department of Medicine, Dartmouth-Hitchcock Medical Center, Lebanon, NH 03766, USA

**Author notes:** Correspondence should be addressed: Chilakamarti V. Ramana.

**Keywords:** Signal transduction pathway, Protein-protein interaction, Transcription factor, Protein kinase, Signaling hubs

## Abstract

Growth factors and cytokines activate signal transduction pathways and regulate gene expression in eukaryotes. Intracellular domains of activated receptors recruit several protein kinases as well as transcription factors that serve as platforms or hubs for the assembly of multi-protein complexes. The signaling hubs involved in a related biologic function often share common interaction proteins and target genes. This functional connectivity suggests that a pairwise comparison of protein interaction partners of signaling hubs and network analysis of common partners and their expression analysis might lead to the identification of critical nodes in cellular signaling. A pairwise comparison of signaling hubs across several related pathways might also reveal novel signaling modules. Analysis of <u>P</u>rotein <u>I</u>nteraction <u>C</u>onnectome by <u>V</u>enn (PIC-VENN) of transcription factors STAT1, STAT3, NFKB1, RELA, FOS and JUN, and their common interaction network suggested that BRCA1 and TSC22D3 function as critical nodes in immune responses by connecting the signaling nodes into signaling modules. Mutations or differential expression levels of these critical nodes in pathological conditions might deregulate signaling pathways and their target genes involved in inflammation. Biological connectivity emerges from the structural connectivity of interaction networks across several signaling hubs in related pathways. Application of PIC-VENN to several signaling hubs might reveal novel nodes and modules that can be targeted to simultaneously activate or inhibit cell signaling in health and disease.

## 1 Introduction

Signal transduction in mammalian cells is mediated through specific membrane receptors and intracellular kinases to regulate gene expression [1]. Ligand binding activates receptor tyrosine kinases (RTKs) activity by auto-phosphorylation or by serving as a substrate for other tyrosine kinases [2]. Receptor phosphorylation can occur at several sites and induce modifications that stimulates intrinsic enzyme activity or establish docking positions for adaptor proteins that recogniz**e** phosphorylated tyrosine residues. In turn, tyrosine RTKs phosphorylate other substrates, thereby initiating a cascade of downstream signals including activation of Serine/Threonine protein kinases and multiple transcription factors. Several enzyme cascades consisting of mitogen-activated protein kinases (MAPK) and stress-activated protein kinases (SAPK) play critical roles in animal development as well as immune responses to pathogens [3]. Protein kinases, and transcription factors can serve as critical hubs or platforms for the assembly of signaling molecules [1-3]. Precision medicine requires the identification of molecular defects in human genetic diseases at the individual and gene-specific level. Targeting multiple signaling pathways is often required to prevent resistance to chemotherapy in cancer [4]. Identification of defects in cell signaling pathways is essential for understanding the molecular basis of human diseases [1-4]. The complexity of signaling pathways arises from the diverse ways protein interactions connect between signaling molecules to regulate gene expression. Several protein interaction databases such as STRING, BIOGRID, and WIKI-PI were developed [5-7]. Furthermore, differential mRNA levels of signaling molecules in a tissue or cell-specific fashion in normal and pathological conditions were also reported in databases such as Gene Expression across Normal and Tumor tissue (GENT), Renal Gene Expression Database (RGED) and Lung Cell Transcriptome Explorer [8-11]. The main objective of this paper is to gain insights into the mammalian signaling pathways by pairwise comparison of protein interaction partners of critical signaling hubs in multiple related signaling pathways. In addition, analysis across several hubs in related pathways might reveal critical nodes and modules involved in cell signaling. It is clear from protein interaction databases that co-operative transcription factors generally share a large number of common protein interaction partners (5-7). Furthermore, these transcription factors also share a large number of common target genes. Identification of common signaling molecules between two signaling hubs and organizing them into a concise interaction network allows for visualizing the critical connections. Critical nodes and modules across signaling hubs were determined by functional data analysis of candidates in various databases from knockout, knockdown or overexpression studies. This approach might reveal modules of functional connectivity across several signaling hubs in related signaling pathways and provide potential targets to simultaneously activate or inhibit gene expression.

## 2 Methods

All entries of protein interaction partners of signaling hubs in this paper were retrieved from WIKI-PI, a human protein-protein interaction database [7]. It is worth pointing out that protein interaction databases differ in the list of interaction partners for each entry based on criteria like in vitro or in vivo detection and methodologies such as immunoprecipitation, co-expression or yeast 2-hybrid interaction analysis [5-7]. They also differ in the number of model organisms included on their list. Schematic representations of cytokine signaling (Figure 2) were modified from previously published reviews [12-14]. Microarray data for innate immunity model represented by mouse macrophage cell line (RAW 264.7) treated with Toll-like receptor [TLR] ligands and adaptive immunity model represented by mouse lung epithelial cells (MLE-Kd) expressing influenza HA antigen co-cultured with CD8^+^ T cells for 6 hours were used [15.16]. Upstream promoter region (−1000 to +1) of 150 induced genes in RAW264.7 cells by TLR ligands or 50 induced genes in MLE-Kd cells by CD8^+^ T cell recognition were analyzed for transcription factor binding sites using P-SCAN software [17]. Transcription factors considered most significant in innate and adaptive pathogen responses were shown. Target genes for RELA, STAT1, JUN, and BRCA1 transcription factors were obtained from Transcriptional Regulatory Relationships Unravelled by Sentence based Text (TRRUST) database [18].

**Figure 1.**
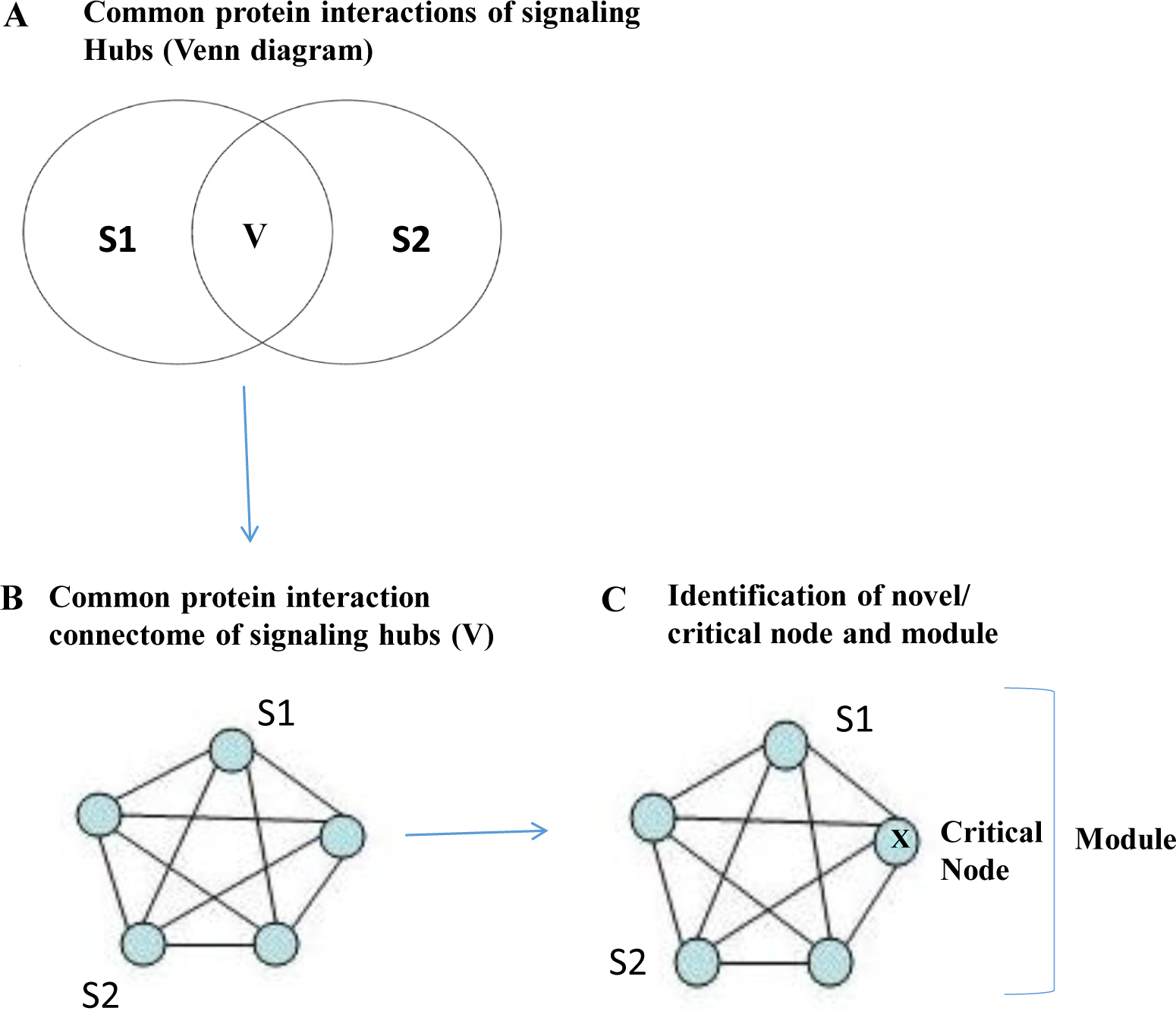
Schematic outline of Protein Interactions Connectome-VENN (PIC-VENN) method. (A) Common protein interaction partners were designated as V for common of signaling hubs S1 and S2, respectively (B) Network pathway representation of common interaction partners. Interacting proteins (nodes) were represented by circles and interactions (edges) by connecting lines. (C) Identification of a critical node and module from the analysis of gene perturbation data of signaling pathway (knockout or overexpression) in databases.

**Figure 2.**
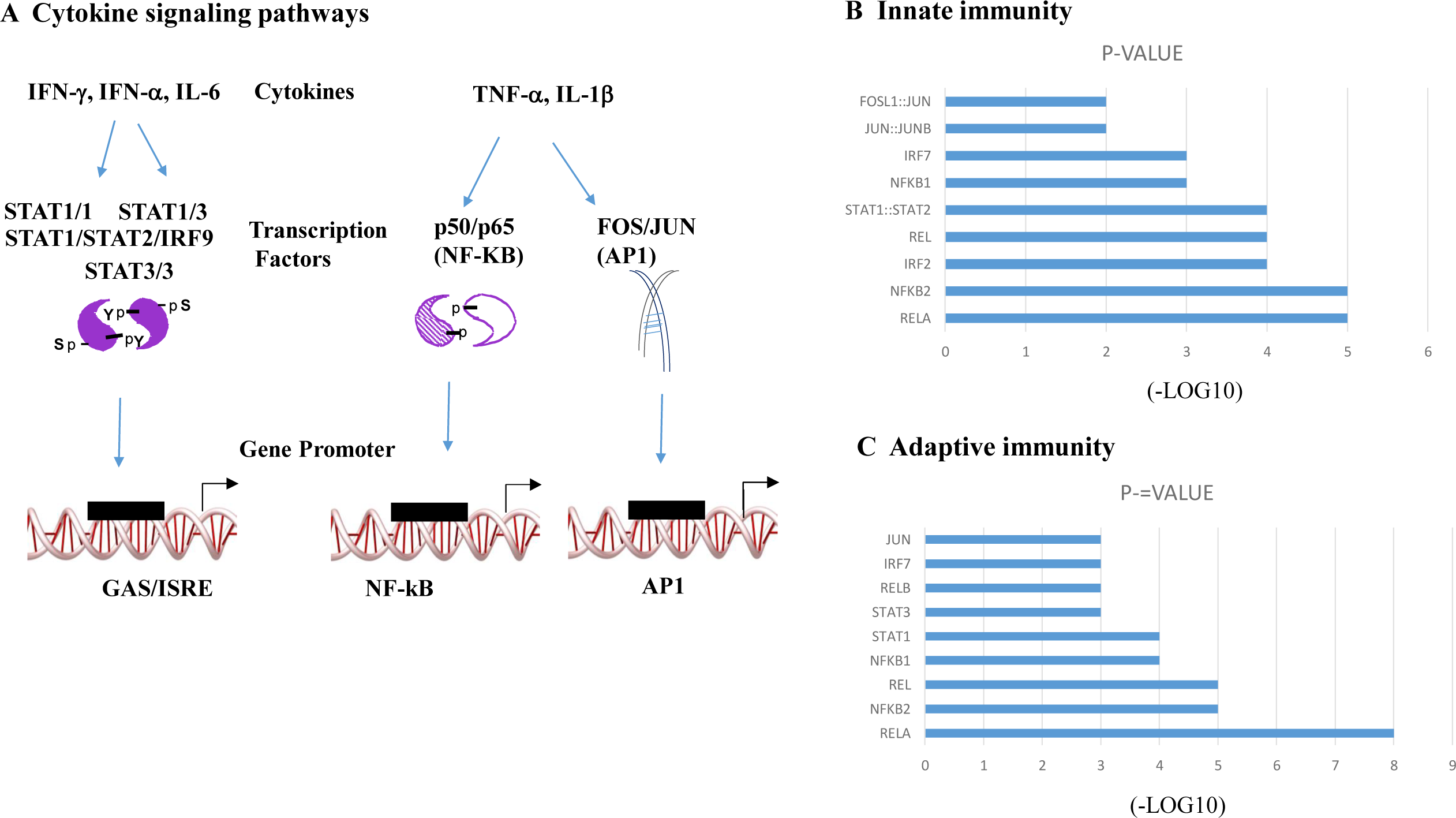
Cytokine signaling and transcription factor hubs critical for innate and adaptive Immunity (A) Schematic representation of cytokines and transcription factors associated with innate and adaptive immunity. (B) Transcription factors in the promoter region (−1000 to +1) of genes involved in the regulation of innate immunity in mouse macrophage RAW 264.7 cells by TLR ligands were ranked by significance. (C) Transcription factors in the promoter region (−1000 to +1) of genes involved in the regulation of adaptive immunity in mouse lung type II cells by CD8^+^ T cell recognition were ranked by significance.

### 2.1 Differential Gene Expression

Tissue expression data for BRCA1, MYC, NOTCH1, and FBXW7 were obtained from Gene Expression across Normal and Tumor tissue (GENT) database [8]. Relative mRNA expression levels of BRCA1 and TSC22D3 in control and dexamethasone treated small airway epithelial cells of human donors were retrieved from the database [11]. Read counts of each gene in the database were obtained by HTSeq and differential expression results were obtained by DESeq2.

### 2.2 Protein Interactions Connectome-Venn (PIC-VENN) method of signaling hubs

In the protein network analysis, interaction partners of signaling hubs S1 and S2 were considered. The overlapping area is V, for “Venn” that includes the elements that are in both sets. This relationship can be expressed in terms of an overlapping set as shown in a Venn diagram (Figure 1). The PIC-VENN method consists of three steps. First, Microsoft Excel software was used to construct a simple Venn diagram of common interaction partners by pairwise comparison of critical signaling hubs. Briefly, protein interaction partner lists of the two signaling hubs were analyzed for duplicate values in the Excel program that correspond to common interaction partners of the signaling hubs. In Step 2, a network diagram of common interaction partners of the hubs together with the critical signaling hubs was visualized in BIOGRID tool Ensign (Human Mine) or the Search Tool for the Retrieval of Interacting Genes/Proteins (STRING) database. The interacting proteins or nodes were represented by circles and-protein interactions by lines or edges in the network. In step 3, novel or critical node was selected from the common interaction partners by determining the biological effect of a node on the signaling pathway from knock out or over expression studies (from National Center for Biotechnology Information/Pubmed query) or from differential expression in a disease-specific gene expression databases such as GENT, Renal Gene Expression Database (RGED) and Lung Cell Transcriptome Explorer (LCTE). Databases such as Gene cards (www.genecards.org) and mouse genome informatics (www.mgi.org) also provide concise information on entries related to human and mouse genes for critical node analysis, respectively.Two or more signaling hubs connected together with a critical node was designated as a signaling module.

## 3 Results and Discussion

### 3.1 STAT, NF-kB and AP1 family of transcription factors as critical hubs in innate and adaptive immunity

Cytokines such as Interferons (IFN), Tumor necrosis factor (TNF-α), Interleukin-1 (IL-1) and Interleukin-6 (IL-6) play an important role in innate immunity [19-21]. Dendritic cells and macrophages of innate immunity patrol various tissues and produce significant amounts of these cytokines in response to virus infection [21]. In contrast, CD8^+^ T cells play an important role in antiviral defense, tumor suppression and in the control of immunopathology in adaptive immunity [22,23]. After engagement of the T cell receptor (TCR) by an appropriate antigen-major histocompatibility complex (MHC) on a target cell, activated CD8^+^ T cells produce pro-inflammatory cytokines such as TNF-α and IFN-γ [23]. These cytokines activate multiple transcription factors that bind to specific DNA sequences or cis-elements to regulate gene expression. Transcriptional regulatory elements in eukaryotes are predominantly located within 1 kilobase [kb] of the transcription start site [24,25]. Intracellular signal transduction pathways play a major role in pathogen response and are regulated by Signal Transducer and Activator of Transcription (STAT), Nuclear Factor-Kappa B (NF-KB) and Activator Protein (AP1) members of transcription factors (Figure 2A). Involvement of JAK kinases, IKB kinases as well as mitogen-activated and stress-activated protein kinases (MAPK/SAPK) such as ERK, JNK, p38 and activation of transcription factors STAT1, NF-KB and AP1 in target cells in response to cytokines released from immune cells has been described [3,26,27]. Diversity of signaling and gene expression are orchestrated by cooperative or alternative binding of distinct members of a transcription factor family. STAT members like STAT1 and STAT3 bind to gamma activated sequence (GAS) as homodimers or heterodimers or as trimeric complex of STAT1, STAT2 and IRF9 to the Interferon stimulated response element (ISRE). AP1 is a leucine zipper domain containing heterodimeric transcription complex comprised of FOS and JUN family of transcription factors [3]. The NF-KB family is a set of five structurally related transcription factors that form homodimers or heterodimers and function as transcriptional regulators. There are two classes of NF-KB family members: class one consists of p50/p105 (NFKB1) and p52/p100 (NFKB2) while class two consists of RELA/p65, RELB and c-REL [27].

The subunit composition of STAT, NF-KB and AP1 transcription factors may vary significantly during inflammatory response. To establish the frequency of the subunit composition of these transcription factors in innate immunity, a dataset of 150 genes regulated by Toll-like ligands (TLR) in RAW 264.7 macrophages were scanned in the promoter region (1 Kb upstream or -1000 to 0) using P-SCAN [17]. Members of NF-KB (RELA, REL, NFKB1, NFKB2); STAT/IRF family (STAT1, STAT2, IRF2, IRF7) and AP1 (FOS, JUN, JUNB) were highly represented (Figure 2B). In addition, a dataset of 50 genes regulated by CD8^+^ T cell recognition of antigen presenting MLE-Kd lung epithelial cells representing adaptive immunity were also scanned in the I kb upstream promoter region. Members of NF-KB (RELA, REL, NFKB2, NFKB1), STAT/IRF family (STAT1, STAT3, IRF7) and AP1 (JUN) were again highly represented (Figure 2C). These results suggested that transcriptional targets and biological functions of innate and adaptive immunity were associated with these three classes of transcription factors. Target genes of signaling hubs of innate and adaptive immunity such as RELA, STAT1 and JUN also share regulation by a common transcription factor network that includes SP1, STAT3, IRF1, FOS, TP53, HIFA, CREB1 and MYC suggesting that functional connectivity is mediated by common protein-protein interactions and target genes (Figure 3).

**Figure 3.**
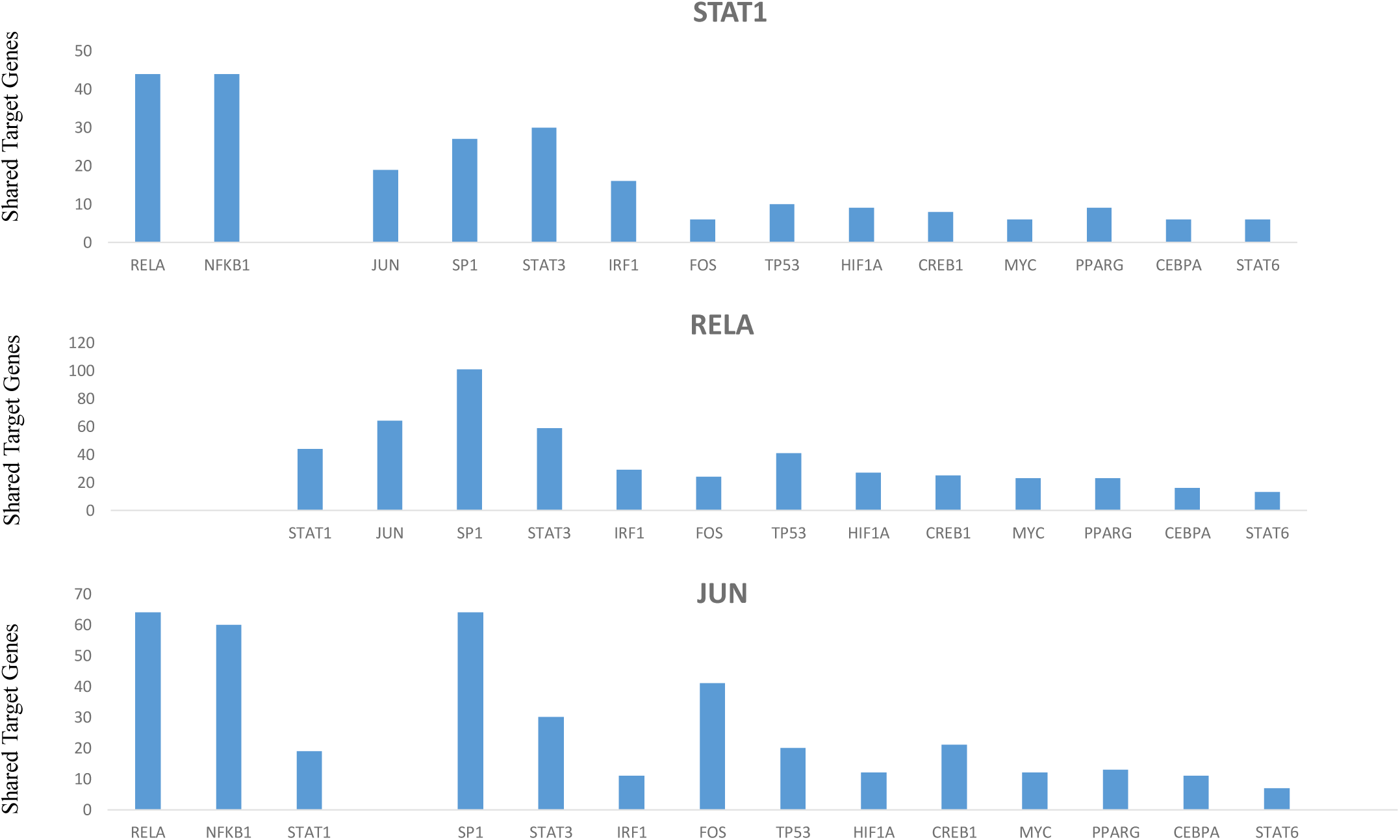
Shared transcription factor profiles in the target genes of STAT1, RELA and JUN.

### 3.2 Case study 1. Application of PIC-VENN to STAT1, STAT3, RELA and JUN hubs

To determine the common interaction partners of these transcription factors, pairwise comparison of critical signaling hubs was performed. Venn diagrams of STAT1 versus RELA, STAT1 versus STAT3 and STAT1 versus JUN indicated that there were 14, 13 and 12 common interactions between the hubs, respectively. Furthermore, comparison of common interacting partners across the hubs revealed that Breast Cancer Gene or BRCA1 emerged as one of the critical components of these signaling hubs (Figure 4A). Network analysis using BIOGRID program eSYN showed that BRCA1 emerged as a node linking RELA, STAT1, and JUN. Protein interactions identified by genetic analysis in the network were shown in green lines and BRCA1 node was highlighted in yellow color. In addition, a module consisting of STAT1, BRCA1, JUN and RELA can be identified across the hubs in pathways (Figure 4B). PIC-VENN results of common interaction partners can be used to search for biological significance in disease-specific expression array databases. Mutations in BRCA1 are common in breast and ovarian cancers [28,29]. Comparison of BRCA1 expression in normal and tumor tissues revealed differential expression in cancers, especially blood cancers (Figure 5A). In addition, cancer cell lines of the blood including lymphoma and myeloma express high level of BRCA1 (data not shown). These results suggest that BRCA1 is a novel and critical node and mutations as well as abnormal level of BRCA1 expression in cells of immune origin may play an important role in cancer. BRCAI is a multifunctional protein with critical roles in cell cycle, DNA repair and tumor suppression [28]. BRCA1 target genes include several transcription factors such as STAT1 and STAT3 involved in immunity as well as TP53, MYC and HIF1A involved in cell growth suggesting that BRCA1 regulates multiple feedback loops that are involved in maintaining cellular homeostasis (Table 1). The target genes of BRCA1 also included IRF7 that plays an important role in interferon signaling as well as production of type I interferon [30]. BRCA1 functions in the transcription control of gene expression by interacting with specific transcription factors like TP53, ER alpha and MYC [31]. BRCA1 also interacts with RNA polymerase holoenzyme complex, histone acetyl transferases such as CBP/p300 and histone deacetylases such as HDAC that regulate nucleosome assembly and disassembly [31,32]. In addition, BRCA1 interacts with DNA methyl transferases and functions as E3 ubiquitin ligase to regulate epigenetic mechanisms such as methylation and ubiquitination [31]. PIC-VENN studies suggest that BRCA1 may have a major role in modulating cytokine signaling and a critical role in innate and adaptive immunity. BRCA1 modulates IFN-γ signaling by specifically binding to STAT1 after phosphorylation on serine 727 residue [33]. This serine phosphorylation of STAT1 was highly enhanced in response to synergistic combination of IFN-γ with a variety of cytokines such as TNF-α and IL-1β as well as bacterial cell wall component LPS induced inflammatory gene expression [34, 35]. IFN-γ caused a decrease in cell proliferation due to apoptosis in cells expressing wild type BRCA1 and mediated by an upregulation of type I IFN production [30,36]. BRCA1 was recently identified as a nuclear co-factor to IFI16 in the sensing of foreign DNA and subsequent cytoplasmic inflammasome formation and secretion of antiviral cytokines such as IL-1β and IFN-β [37]. STAT3 and RELA signaling hubs share 20 common interaction partners. In addition, BRCA1 emerged as a critical node for the RELA-STAT3-JUN-STAT1 module (Figure 5B). STAT3 and RELA share about 50 target genes including cytokines, chemokines, matrix metalloproteases and regulators of cell growth and apoptosis (data not shown). Recent studies have shown that cytokines that predominantly activate STAT3 and RELA such as IL-6 and TNF-α in combination promote differentiation of human monocytic myeloid-derived suppressor cells (MDSC) into inflammatory macrophages [38]. These results are consistent with a role for BRCA1 in inflammation, in addition to its well-recognized roles in transcription, cell cycle and DNA repair [32]. Furthermore, BRCA1 also emerged as a critical node in a pairwise comparison of RELA vs JUN or STAT3 vs JUN signaling hubs (data not shown). Treatment with dexamethasone, a synthetic glucocorticoid receptor agonist suppressed BRCA1 mRNA levels in human airway epithelial cells (Figure 5C).

**Table 1.** BRCA1 target genes common between RELA, STAT1, STAT3 and JUN-regulated genes [highlighted in pink].

| RELA | STAT1 | STAT3 | JUN |
| --- | --- | --- | --- |
| CCND1 |  | CCND1 | CCND1 |
| CDKN1A | CDKN1A | CDKN1A | CDKN1A |
| CYP19A1 |  | CYP19A1 | CYP19A1 |
|  |  | DDIT3 | DDIT3 |
| EGFR | EGFR | EGFR | EGFR |
| MYC |  | MYC | MYC |
| VEGFA |  | VEGFA | VEGFA |
| HIF1A |  | HIF1A |  |
| IFNA1 | IFNA1 |  |  |
| IRF7 | IRF7 |  |  |
| TFF1 |  |  | TFF1 |
| STAT1 |  | STAT1 |  |
| STAT3 | STAT3 |  |  |
|  | JAK2 | JAK2 |  |
|  | KRT17 | KRT17 |  |
| CXCL1 | STAT2 | TP53 |  |
| CYP1A1 |  | MDC1 |  |
| SIRT1 |  | FOS |  |
| EGR1 |  | CDKN1B |  |
| BRCA2 |  |  |  |
| CCNB1 |  |  |  |

**Figure 4.**
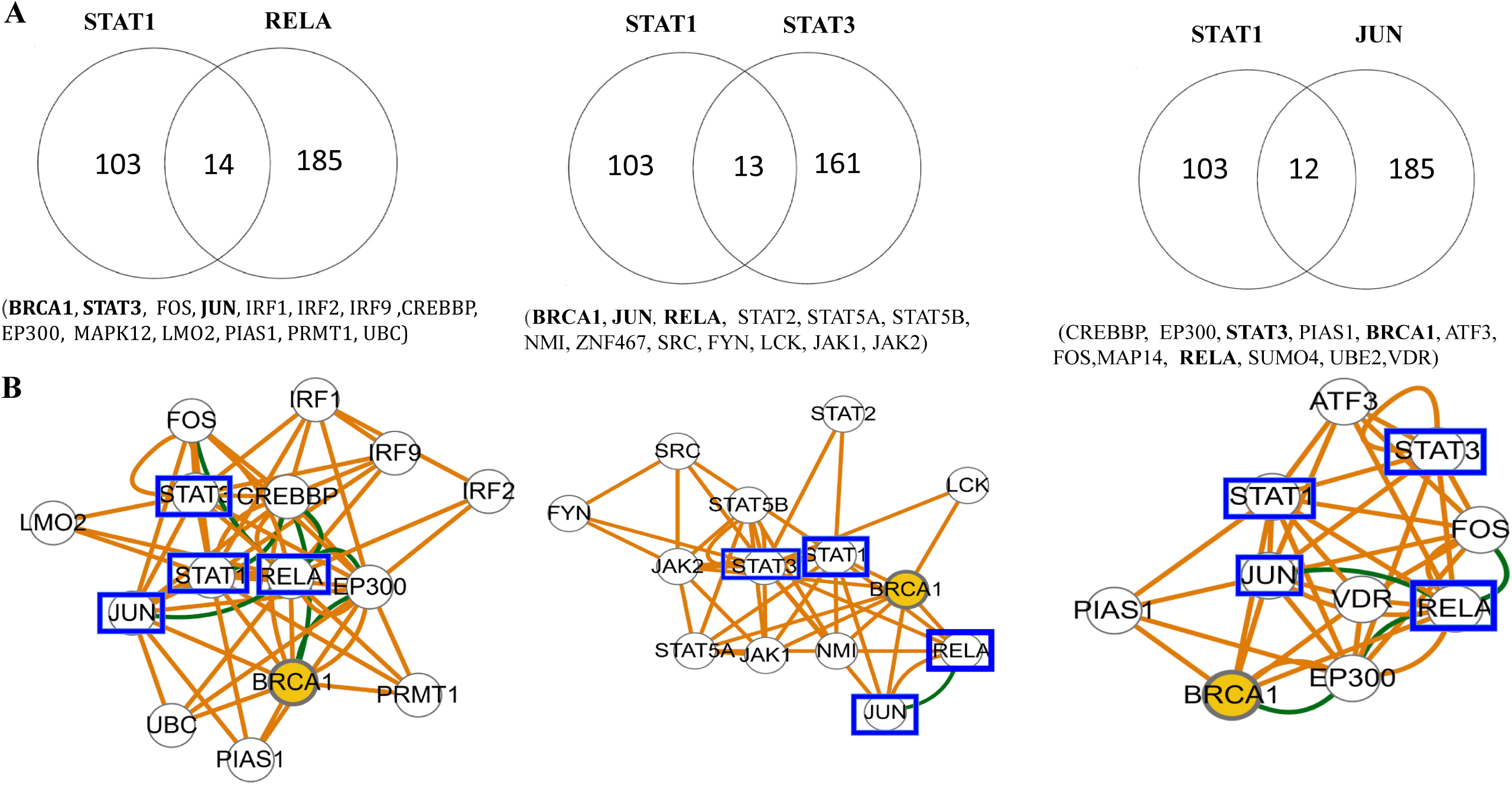
Representation of common protein interaction partners by pairwise comparison of STAT1 versus RELA, STAT1 versus STAT3 and STAT1 versus JUN. (A) Venn diagram (B) Network analysis of common protein interaction partners using eSYN program in BIOGRID database. Critical node (BRCA1) was highlighted in yellow color, components of the module were indicated by blue boxes and genetic interactions by green lines.

**Figure 5.**
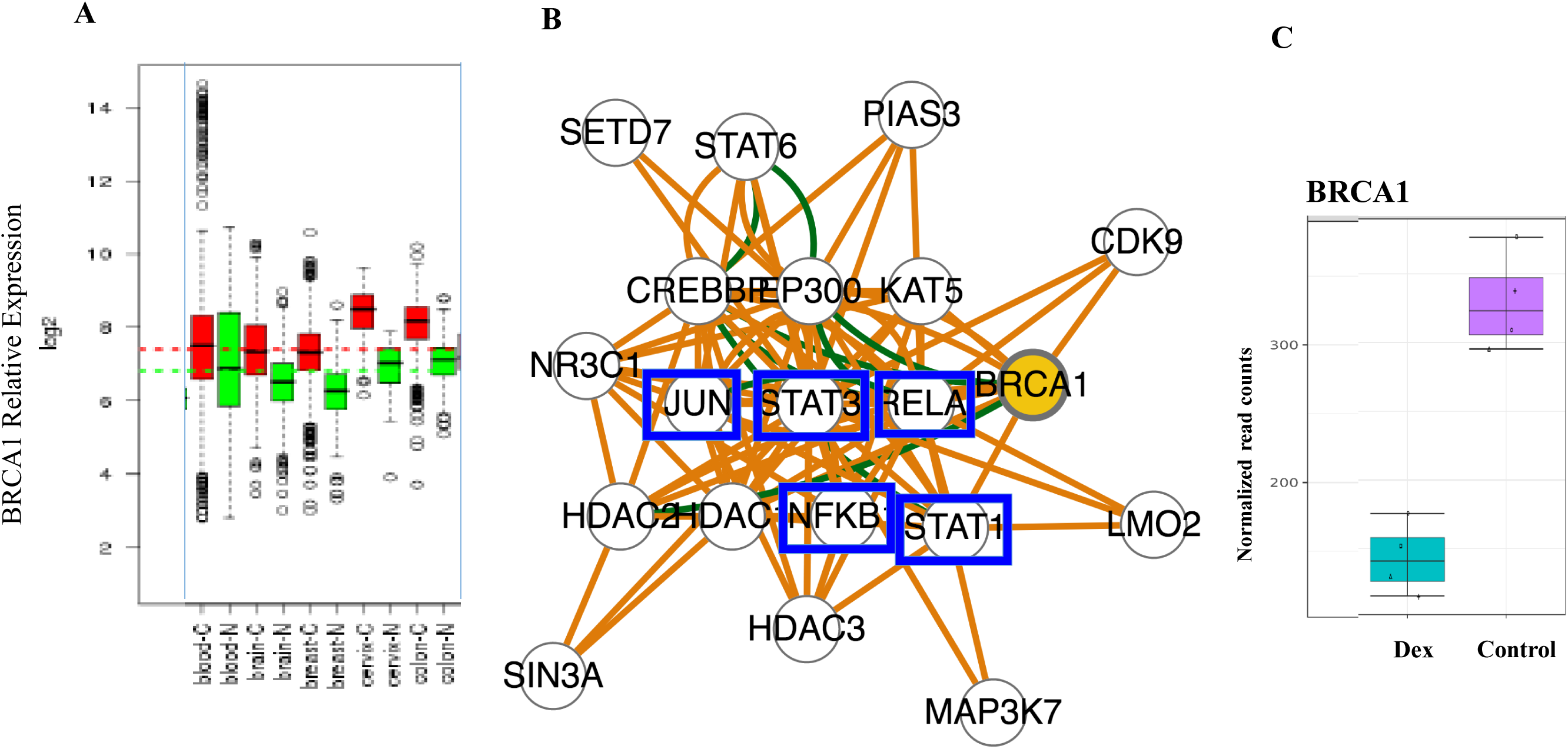
Relative mRNA expression levels of BRCA1 in normal and cancer tissues and BRCA1 as a novel/critical node for RELA and STAT3 hubs. (A) Box plots in selected tissues include blood, brain, breast, cervix and colon. Cancer (C)\ and Normal (N) tissues were indicated by Red and Green, respectively. (B) Network analysis of common interaction partners of RELA and STAT3 signaling hubs. Critical node (BRCA1) was highlighted in yellow color, components of the module were indicated by blue boxes and genetic interactions by green lines. (C) Down-regulation of BRCA1 mRNA expression by dexamethasone in airway smooth muscle cells. Data from eight human donors were retrieved from Lung Cell Transcriptome Explorer. Control and Dex refer to cells untreated or treated with dexamethasone.

IFN-γ is involved in the amplification of the inflammatory response by synergy in signaling with endotoxin, TNF–α and IL-1β and this synergy further enhances the severity of inflammatory response in sepsis [23,34,35]. Collaboration of multiple cis-elements and altered dimer composition of these three classes of transcription factors leading to cross-talk between signal transduction pathways may play an important role in the diversity of gene expression by transcriptional synergy or antagonism.

### 3.3 Case study 2. Application of PIC-VENN to AP1 and NF-KB signaling hubs

In addition to inflammatory response, members of NF-KB and AP1 transcription factors are also activated by diverse stimuli such as cell growth, DNA damage and apoptosis [3,27]. A pairwise comparison of interaction partners of signaling hubs (FOS versus RELA, JUN versus NFKB1 and FOS versus JUN) revealed that these three sets also share a small number of interaction partners within the hubs and across the hubs (Table 2). Network visualization of the common interaction partners across two (NFKB1 versus JUN and FOS versus JUN) or all three (NFKB1 versus JUN, FOS versus RELA and FOS versus JUN) pairs of hubs suggested that TSC22D3 or glucocorticoid-inducible leucine zipper (GILZ) may act as a novel and critical node linking the module FOS, JUN and NFKB1 (Figure 6A and 6B). Consistent with this prediction, functional studies have revealed that TSC22D3 interacts with NF-KB and AP1 and negatively regulated its function [39,40]. AP1 and NF-KB member connections to negative regulators of signaling such as BCL3 and ATF3 were also highlighted (Figure 6A). Interestingly, dexamethasone treatment significantly induced TSC22D3 mRNA levels in human airway epithelial cells (Figure 6C).

**Table 2.** Shared common interaction partners across NF-KB and AP1 Signaling hubs.

|  |  |  |
| --- | --- | --- |
| NFKB1 | FOS | FOS |
| VS JUN | VS RELA | VS JUN |
| AR | CARM1 | ATF2 |
| ATF3 |  | ATF3 |
| BCL3 |  | BCL3 |
| BRCA1 | CREBBP | CREBBP |
| COPA5 | EEF1D | BATF |
| ELF3 | JUN | ATF4 |
| EP300 | KDM2A | CEBPE |
| ESR1 | MAP3K7 | CEBPG |
|  | NCOA3 | NCOA3 |
| FOS |  | CREB5 |
| GSK3B |  | COPS4 |
| HMGA1 | CSNK2A1 | CSNK2A1 |
| IKKBK | CSNK2A2 | CSNK2A2 |
| NCOA1 |  | NCOA1 |
| NCOA6 | NCOA6 | NCOA6 |
| NCOR2 | NCOR2 | NCOR2 |
| NR3C1 | PML | GTF2 |
| PPARG | PRKACA | MAF |
| PPP4C | SIRT1 | MAFB |
|  | NFKB1 | MAPK1 |
| SP1 | UBE2D1 | NACA |
|  | UBE2L3 | ETS2 |
| STAT3 | USF2 | NCOA2 |
| TRIP4 |  | DDIT3 |
| ETS1 |  | ETS1 |
|  |  | APP |
|  |  | NELFB |
|  |  | RB1 |
| RELA |  | RELA |
|  |  | RPS6KA2 |
|  |  | RUNX1,2 |
|  | SMAD3 | SMAD3 |
|  |  | SMARCD1 |
| SPI1 |  | SPI1 |
|  | STAT1 | STAT1 |
|  |  | SUMO1,2,3,4 |
|  | TBP | TBP |
|  | TAF1 | TAF1 |
| TSC22D3 | TSC22D3 | TSC22D3 |
|  |  | UBE2L |
|  |  | VDR |

**Figure 6.**
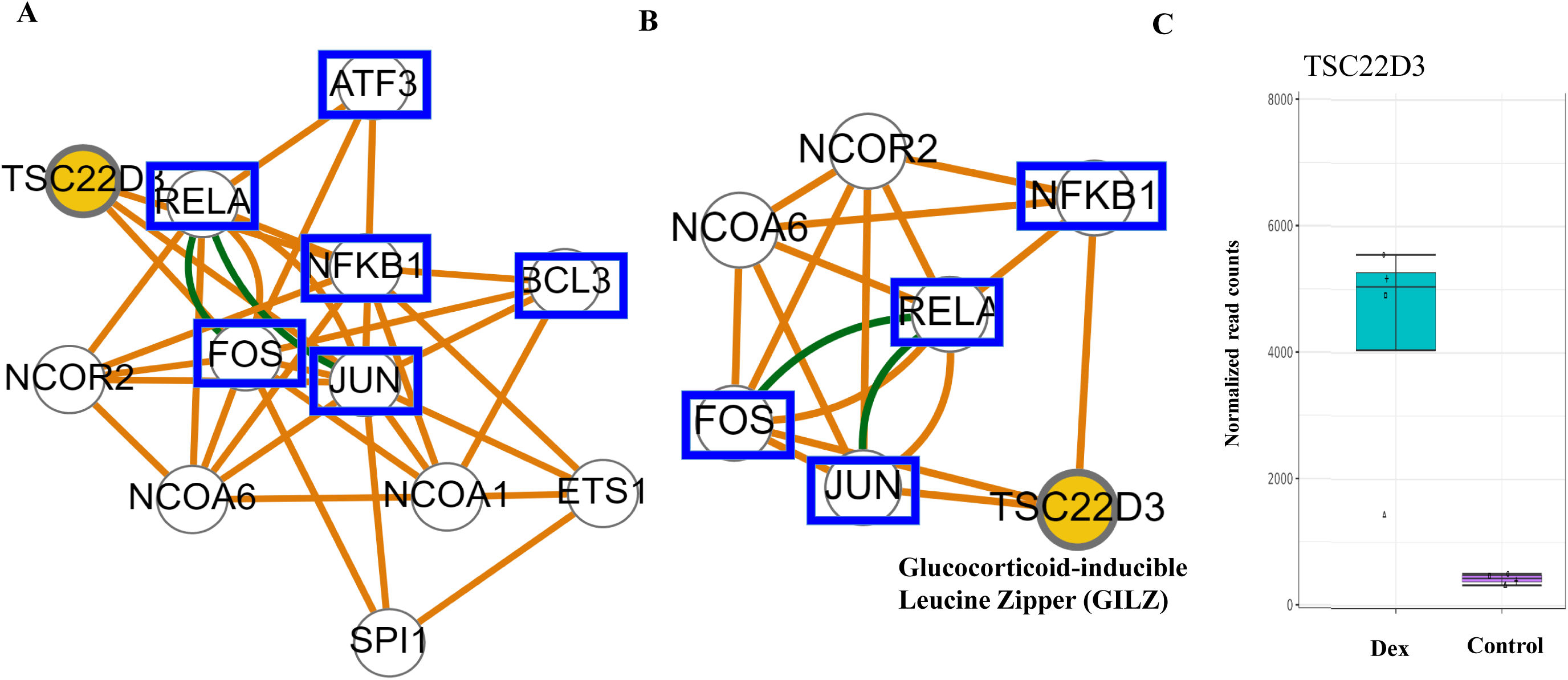
Network analysis of common interaction partners across NFKB and AP1 hubs. (A) Signaling pathway across two pairs of hubs (NF-KB1 versus JUN and FOS versus JUN) (B) Signaling pathway across three pairs of hubs (NFKB1 versus JUN, FOS versus RELA and FOS versus JUN). Novel/critical node (TSC22D3 or GILZ) was highlighted in yellow color, components of the module were indicated by blue boxes and genetic interactions by green lines. (C) Up-regulation of TSC22D3 mRNA expression by dexamethasone in airway smooth muscle cells. Data from eight human donors were retrieved from Lung Cell Transcriptome Explorer. Control and Dex refer to cells untreated or treated with dexamethasone.

### 3.4 Case study 3. Application of PIC-VENN to JAK Kinases in Interferon Signaling

JAK Kinases are critical signaling hubs in growth factor and cytokine signaling including interferon responses [13,26]. Virus infected cells produce different forms of interferons that serves as a danger signal to the neighboring cells to establish an antiviral response [20]. Dendritic cells and macrophages produce type I Interferon (IFN-α/β) after virus infection [19-21]. In contrast, activated NK cells and CD8^+^ T cells produce type II Interferon or IFN-γ [22,23]. JAK1 and TYK2 are activated in type I Interferon signaling while JAK1 and JAK2 are activated in type II Interferon signaling [26]. Interferon binds to a specific and distinct receptor on the cell surface and activate JAK-STAT pathway. This pathway is involved in the transcriptional regulation of several hundred genes [13]. Several Stat1-independent pathways are also activated simultaneously, including extra-cellular signal regulated kinase (ERK1/2), SAPK members such as P38, phosphatidylinositol-3 kinase (PI-3K) and SRC kinases [41,42]. The role of JAK kinases in tyrosine phosphorylation and creation of docking site and activation of STAT signaling is well established [13]. The role of additional tyrosine phosphorylation sites on the IFN receptor and JAK Kinase interaction partners involved in alternative signaling pathways might play a role in Stat1-independent IFN signaling remains to be established [41,42]. However, limited information is available on JAK kinase interacting protein partners involved in these alternative pathways. Application of PIC-VENN revealed that JAK1 and TYK2 share 16 common interaction partners and JAK1 and JAK2 share 44 interaction partners. In addition, type I and type II IFN signaling share 11 common interaction partners (Table 3). JAK1 and JAK2 common interaction network is large and visualized into two separate sub-networks with roles in IFN-γ and growth factor signaling (Figure 7A and 7B). Common network interaction partners highlight the role of JAK1 and JAK2 in STAT1 and STAT3 activation as well as P-I-3-kinase and RAF1 involvement in AKT1 and ERK1/2 activation, respectively [41]. Analysis of protein interaction network suggested a novel connecting module of P-I-3-kinase subunit (PI3KR1) to Phospholipase gamma (PLCG) and Src homology 2 (SH2) domain containing adaptor (SHB/SH2B2) and Signal Transducer and Adaptor (STAM) to JAK2 in IFN and growth factor signaling. PLCG is involved in the generation of second messengers diacyl glycerol (DAG) and inositol 1,4,5-trisphosphate (IP3). DAG leads to the activation of protein kinase C theta and IP3 imediates an increase in the intracellular concentration of calcium (Ca^2+^) and activation of the Ca^2+^-dependent kinases and phosphatases [43]. In addition, PLCG plays an important role in lung epithelial regulation of cell adhesion molecule (ICAM1) by IFN-γ [44]. The SHB adaptor (SHB) levels were regulated by interferon, and plays a role in apoptosis and tumor suppression [45-47]. Thus, PIC-VENN method is capable of revealing novel interacting partners between JAK Kinase signaling hubs in Interferon signaling. Furthermore, this analysis suggests several ways to test the connection between specific protein interaction and activation of novel components (highlighted by orange boxes in figures) of the signal transduction pathways. IFN-α/β as well as IFN-γ activate P-I-3-kinase and AKT1 signaling [48,49]. JAK1 and TYK2 phosphorylate and activate IRS1/IRS2 which in turn activate PI3K in IFN-α/β signaling [50]. This pathway is evident in the common interaction network of IFN α/β signaling (Figure 7C). However, JAK1 and JAK2 are not involved in IRS1/IRS2 activation in IFN-γ signaling [50,51]. Protein interaction databases differ in the lists of protein interaction partners of each entry leading to novel pathways. Visualization of the common interactions between the signaling hubs and across the signaling hubs in to a pathway in different databases helps to appreciate the similarities and differences between the signal transduction pathways. Therefore, network visualization of common interaction pathways in different databases might be useful to gain novel insights into signaling networks.

**Table 3.** Common interaction partners of JAK kinases in Interferon signaling.

| Signaling Pathway | Hub1 and Interactions | Hub2 and Interactions | Common Interaction Partners |
| --- | --- | --- | --- |
| IFN $\gamma$ | JAK1 (69) | JAK2 (94) | 44 (IL3RA, IL5RA, IL4R, INSR, **IRS1**, **IRS2**, IFNGR1,**GHR**, FES, BRCA1 ABL1, ELP2, ERBB2, CXCR4, CCR5, CSF3R CSF2RB, JAK3, MDK, OSMR, **PDGFRB**, **PIK3R1**, PLCG2, **PRMT5**, **PTPN6**,PTPN11, PTK2B,PTPRC RAF1, SH2B2, SOCS1, SOCS3, STAM1, **STAM2**, GRB2, HIST3H3, IGF1R, ,TNFRSF1A, **STAT1**, **STAT2**, STAT3, STAT5A, TEC, TSHR) |
| IFN $\alpha/\beta$ | JAK1 (69) | TYK2 (36) | 16 (EIF2AK2, GHR, GNB2L1, IFNAR2, **IRS1**, **IRS2**,JAKMIP1 **PDGFRB**, **PI3KR1**, PLAUR, **PRMT5**,**PTPN6**,PTPRC, **STAM2** **STAT1**, **STAT2**) |

**Figure 7.**
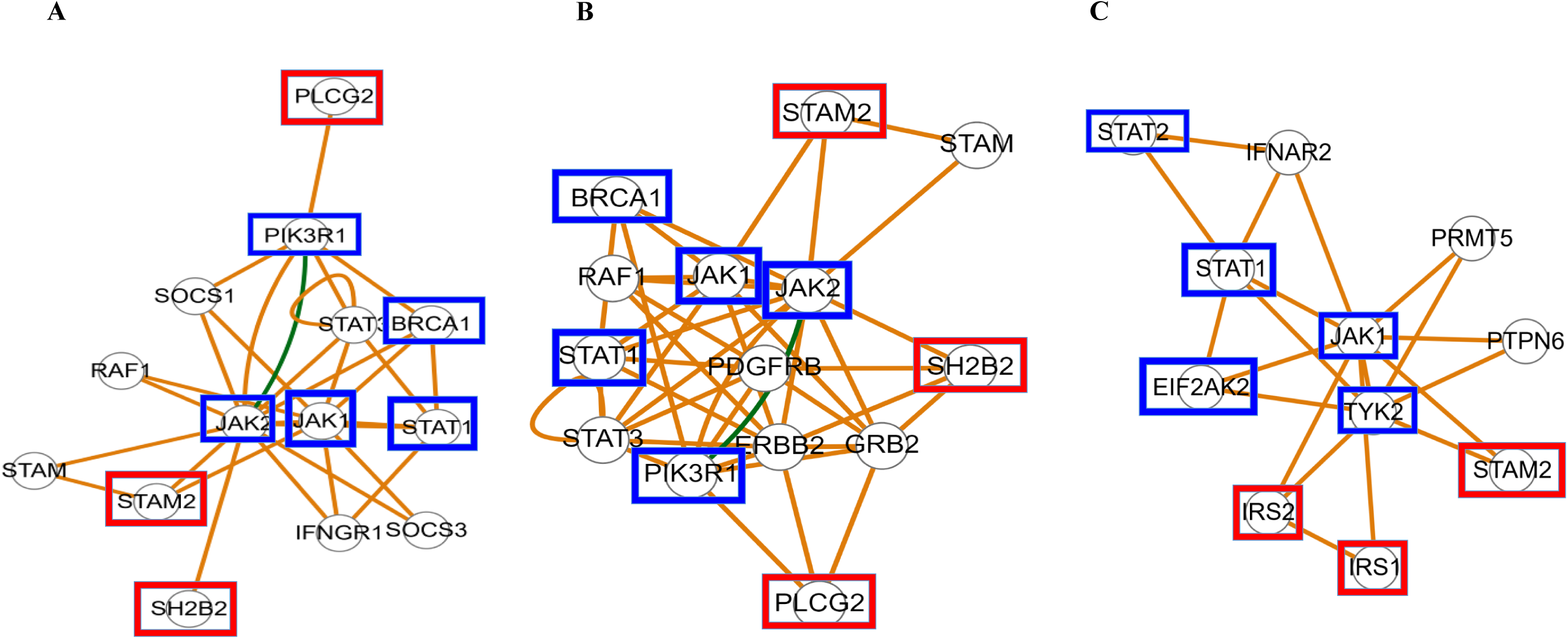
Common protein interactions of JAK kinases and their network analysis in interferon and growth factor signaling (A) Common protein interactions of signaling hubs JAK1 and JAK2 and their network analysis in IFN-γ signaling (B) Common protein interactions of signaling hubs JAK1 and JAK2 and their network analysis in growth factor signaling (C) Common protein interactions of signaling hubs JAK1 and TYK2 and their network analysis in IFN α/β signaling. Components of the module were indicated by blue boxes. Novel nodes identified in this study were indicated by orange boxes.

### 3.5 Case 4. Application of PIC-VENN for elucidation of signal transduction pathways and potential targets involved in disease conditions

Sepsis and septic shock represent the pathological response to infection [14]. Cytokines such as Interferon-γ and TNF-α as well as double-stranded RNA and bacterial LPS activate STAT1 and RNA-dependent protein kinase (EIF2AK2 or PKR) signaling hubs in septic shock resulting in inflammatory response, apoptosis and tissue injury [52-54]. Apoptosis plays an important role in sepsis-induced mortality [53,54]. One of the major control points in apoptosis is the sequential activation of Caspases [54]. Studies have indicated that serum from septic shock patients activated transcription factors such as STAT1, NF-kB and IRF1 as well as Caspases [55]. Pairwise comparison of STAT1 and EIF2AK2 signaling hubs interactions revealed that common interacting partners include CASPASE 3, CASPASE 7, STAT3, RAC1 and TYK2 suggesting that caspase activation and apoptosis are an important part of the acute inflammatory response (Table 4). STAT1 regulates the constitutive levels of Caspase expression [56]. JAK kinases such as TYK2 were shown to be required for the activation of STAT1 and EIF2AK2 in response to double-stranded RNA or Bacterial RNA [57]. Inhibition of EIF2AK2 attenuated inflammatory response, blocked apoptosis and improved mortality in a mouse model of acute lung injury [53]. The network diagram of common interaction partners highlighted these experimentally validated results (Figure 8A). Interaction network visualization also suggested that RAC1 and STAT3 interaction might be important for critical nodes STAT1 and EIF2AK2 to form a module involved in the regulation of inflammatory response and disease [58,59]. Acute inflammation is a hallmark of high mortality in pathogen response while chronic inflammation is a characteristic feature of many inflammatory diseases including cancer [60]. Cancer development involves alteration of signal transduction pathways leading to constitutive activation of oncogenes or loss of tumor suppressors implicated in the cell cycle regulation. In addition, several studies have shown that two hits in these critical genes are required to initiate cancer [4,61]. All cancers share some molecular features referred to as hallmarks such as self-sufficiency in growth signals, loss of response to inhibitory growth signals, ability to evade programmed cell death or apoptosis and unlimited replication potential [62]. Molecular defects in cancer are heterogeneous. It is important to understand the critical signaling hubs and interaction partners perturbed in individual signaling pathways in order to achieve the goals of personalized medicine [1,2,4], Targeting common interaction partners of two signaling hubs in a signal transduction pathway may enhance chemotherapy in cancer (Table 4). The following examples include using PIC-VENN to target transcription factors or kinases or both in various cancers. STAT3 and AKT1 constitute potent survival signals for cancers and are constitutively activated in a variety of cancers [63]. These two signaling hubs share 9 interaction partners including the module containing critical nodes SRC and MTOR Kinases (Figure 8B). Furthermore, inhibition of these Kinases can be used in combination to target both pathways in some cancers [64]. Two Serine Kinases STK39 and AKT1 share the same interaction Protein Kinase, p38 or MAPK14 (Table 4). Consistent with a critical function of MAPK14 in cancer, inhibition of the STK39 by antisense approach stabilized microtubules, inhibited cell cycle progression and induced apoptosis. This effect was mediated by inhibition of MAPK14 phosphorylation and enhanced paclitaxel sensitivity of ovarian cancer [65]. Constitutive activation of transcription factors MYC or NOTCH1 occurs in a variety of cancers [66]. These two signaling hubs share 6 interaction partners, including FBXW7 (Figure 8C). Comparing the expression levels of two signaling hubs and a common interaction partner in the normal and cancer tissue database provided clues on the role of this pathway in a particular type of cancer (Figure 9). MYC and NOTCH1 as well as common interactor FBXW7 expression levels were significantly altered in several cancers, especially in blood cancers [67]. Consistent with the results, mutations in FBXW7 were reported in 2-6% of chronic lymphocytic leukemia (CLL) resulting in the activation of NOTCH11 intracellular domain (NCID) and MYC target genes [68]. Network visualization also revealed that RELA, GSK3B components of the module might regulate these critical developmentally regulated genes (Figure 8C). Negative results such as lack of common interaction partners between two signaling hubs can also be informative suggesting that inhibitors for both hubs can be combined into a single chemotherapy.

**Table 4.** Common interaction partners of signaling hubs involved in pathological conditions.

| Signaling Pathway | Hub1 and Interactions | Hub2 and Interactions | Common Interaction Partners |
| --- | --- | --- | --- |
| Septic shock |  |  |  |
| DS RNA | STAT1 (103) | EIF2AK2(48) | 5 (CASP3,CASP7,RAC1,STAT3,TYK2) |
| Bacterial RNA |  |  |  |
| Cancer | STAT3 (161) | AKT1 (191) | 9 (AR,ARFIP2,BRCA1,SRC,RAC1,PTPN1,NR4A1,MTOR,GATA2) |
|  | MYC (126) | NOTCH1(56) | 6 (GSK3B,FBXW7,RELA,KAT2B,SMAD3,YY1) |
|  | AKT1 (191) | STK39 (13) | 2 (MAPK14, WNK4) |

**Figure 8.**
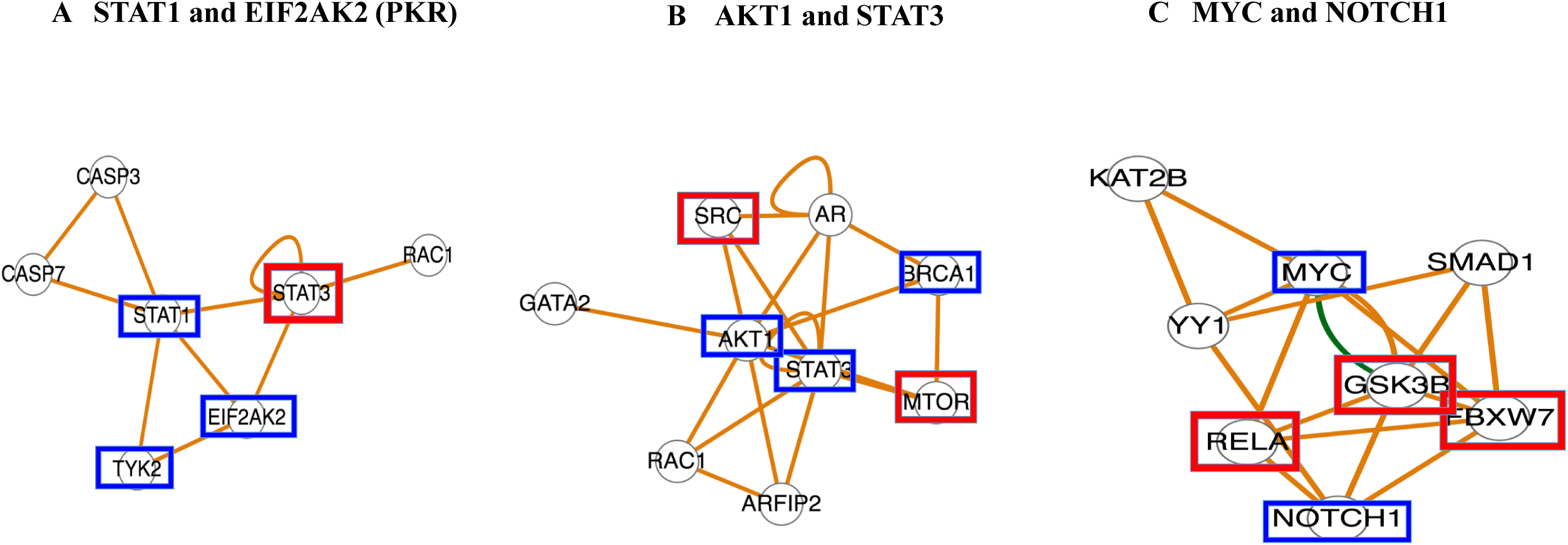
Network analysis of common protein interaction partners of signaling hubs (A) STAT1 versus EIF2AK2 (B) AKT1 versus STAT3 and (C) MYC versus NOTCH1 using eSYN program of Biogrid database. Components of the module were indicated by blue boxes. Novel nodes identified in this study were indicated by orange boxes.

**Figure 9.**
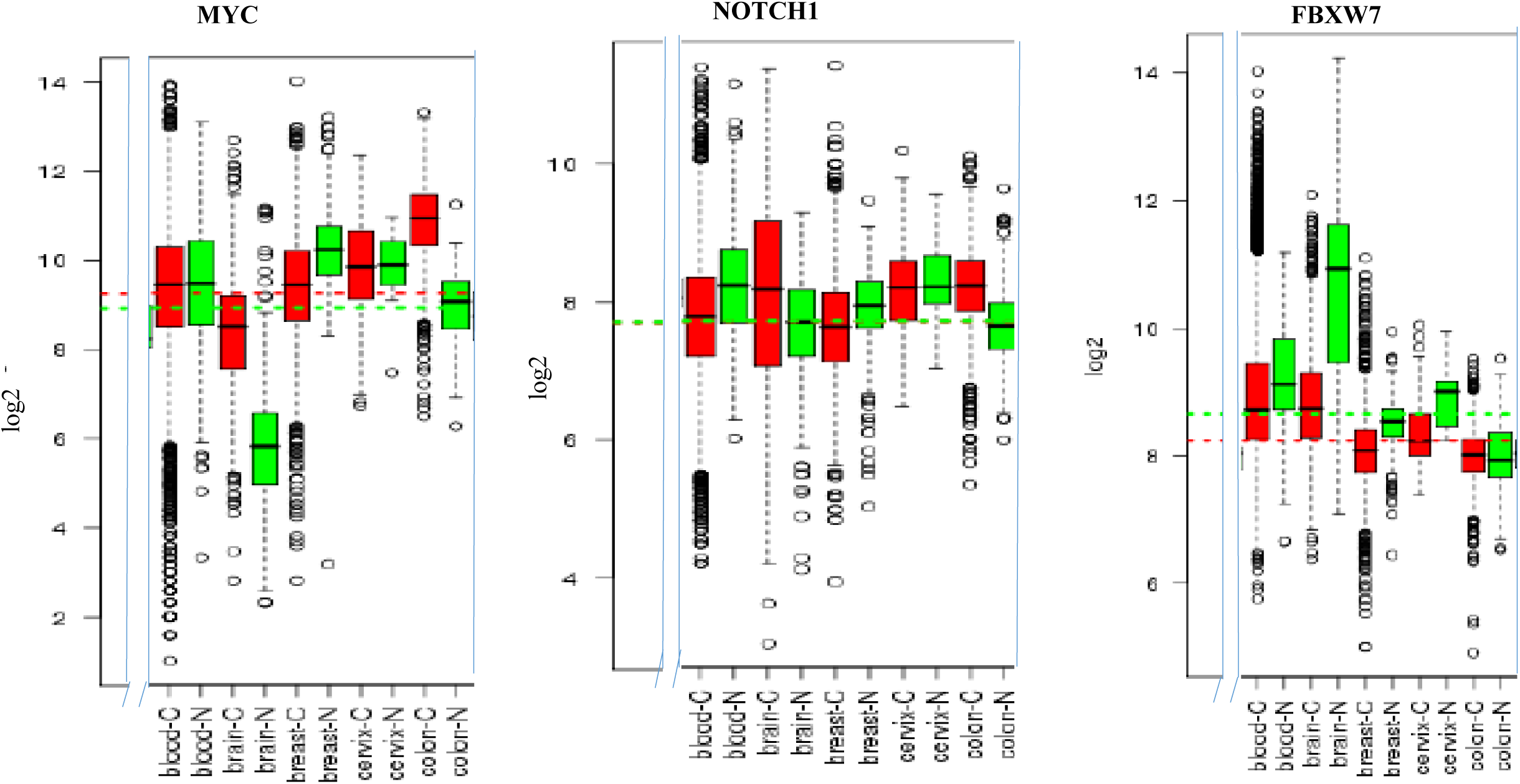
Relative mRNA expression levels of MYC, NOTCH1 and FBXW7 in normal and cancer tissues. Box plots in selected tissues including blood, brain, breast, cervix and colon were shown. Selected regions of the figures were shown for comparison. Cancer (C) and Normal (N) tissues were indicated by Red and Green, respectively.

## 4 Conclusions and Perspectives

In summary, signaling hubs in simultaneously activated signaling pathways share a large number of common interaction partners and target genes involved in a common biological function suggesting that structural connectivity translates into functional connectivity of common interaction partners. PIC-VENN provides significant information on critical nodes and modules when the analysis is carried out across 3 or more pairs of signaling hubs in related signaling pathways (Figures 4 and Figure 6). Even a comparison of two critical signaling hubs provides some useful information on novel nodes and modules (Figure 8). Algorithms such as Google PageRank and Eigenvector Centrality might be helpful in the prediction and ranking of critical nodes from the analysis of gene perturbation data in the future. Small molecules such as dexamethasone can be used to target critical nodes such as BRCA1 and TSC22D3 to simultaneously activate or inhibit multiple target genes. The common protein interaction connectome analysis of critical signaling hubs by PIC-Venn approach is conceptually similar to the identification of shared genes and signal transduction pathways in a comparative study of multiple sclerosis and in response to interferon beta treatment [69]. However, this method was applied to genome-wide analysis (GWAS) of a disease and not critical signaling hubs as detailed in this report. Human protein interaction miner (HPIminer) is a protein interaction database that allows combined view of common and unique interacting proteins of signaling hubs in a Cytoscape program [70]. The entries were curated from Human Protein Reference Database (HPRD). Direct as well as indirect interactions between signaling hubs were also displayed. However, it is difficult to follow or identify unique patterns between hubs or across several hubs when large number of common interactions are involved. Furthermore, the database is limited to human proteins and does not allow for custom building of selective common interactions into a network. In contrast, PIC-VENN method can be used for multiple species and elucidation of novel or critical nodes and modules. A major disadvantage of many protein interactome studies is that they provide a snapshot of expression profile in cells or tissues at a particular time. However, it is well known that protein interactions and gene expression are dynamically and temporally regulated during cellular signaling. Further studies are required to systematically perturb protein interaction network to understand the role of individual nodes and modules in the dynamic nature of signal transduction pathways [71]. The impact of microbiome and its products such as small molecules and small proteins on the regulation of mammalian signal transduction pathways also remains to be investigated [72].

## Acknowledgements

I would like to acknowledge the technical support from the Microarray and Core facilities at Yale University (New Haven, CT) and Dartmouth-Hitchcock Medical Center (Lebanon, NH).

